# The Adjuvant Effects of Amphipathic Helical Peptides

**DOI:** 10.1101/2023.03.13.532490

**Authors:** Jyun-Hong Lyu, Gunn-Guang Liou, May Wang, Ming-Chung Kan

## Abstract

Subunit vaccines are the focus of research in developing new vaccines against infectious diseases. Due to the low immunogenicity of recombinant proteins, adjuvants are required for the activation of humoral and cellular immunity against a protein antigen. In this study, we reported the identification of a novel pathway that can activate humoral immunity against a recombinant protein without inducing inflammatory responses. By fusing an amphipathic helical peptide to GFP, one can increases the immunogenicity of GFP by up to 1000 fold. This enhancement was correlated with the ability of amphipathic helical peptides to bind to cell membranes and cause lysosomal membrane permeabilization. We showed evidence that the amphipathic helical peptide may induce the delivery of antigen across the lysosomal membrane into cytosol. Amphipathic helical peptide fusion provided a new pathway for stimulating immune responses against recombinant proteins.

## Introduction

The challenge of developing a subunit vaccine focuses on two aspects: first, finding the right antigen that may serve as a countertarget for the host immune system; and second, enhancing the specific host immune responses through various mechanisms to a level that can counter the pathogen but will not cause harm to the host. The antigens for subunit vaccines come from various components of the pathogen: polysaccharides, viral proteins that mediate infection, and toxins that cause host morbidity or mortality. These antigen candidates are poor at activating the host immune system and often require the addition of an adjuvant. The adjuvant, according to the FDA, is a compound that is added to or used in conjunction with the vaccine antigen to augment, potentiate, or target a specific immune response to the antigen [1]. The adjuvant may be presented to the immune system in different forms and activate immune responses through various pathways.

The amphipathic peptide is characterized by the propensity of a peptide to form both hydrophilic and hydrophobic surfaces and mediates the interactions between protein and membrane [2]. The M2 amphipathic helical (M2AH) peptide is the membrane anchor of the type A influenza virus proton pump [3]. M2 proteins form a cross-membrane tetramer, transporting protons needed for endosome acidification and viral particle escape [3]. The M2AH resides on the cytoplasmic surface of the host membrane and induces membrane curvature and scission in a cholesterol concentration-dependent manner [4-6]. In a phospholipid bilayer containing cholesterol, the binding of M2AH induces membrane pit formation and a weakening of the lipid bilayer [7]. An M2AH variant is able to induce cholesterol-dependent membrane leakage or membrane lysis [4]. In this study, we have identified several amphipathic helical peptides that stimulated strong humoral immunity when they were fused with a model antigen, green fluorescent protein (GFP) and used in immunization.

## Result

To test the effects of an amphipathic helical peptide on the immunogenicity of a recombinant protein, we fused a peptide (AH2) from the M2 protein of a type A influenza virus strain, A/TW/3355/97(H1N1) to green fluorescent protein (GFP). The fusion proteins were expressed and purified from *E. coli* strain BL21(DE3). The purified fusion proteins were used for the immunization of BALB/c mice using a prime and boost protocol. A variant of the AH2 peptide (AH2M3) that contains three substitutions that disrupted the membrane binding activity of AH2 was used as a negative control. After immunization, the anti-GFP IgG titer was followed for up to 6 months. Compared to His-GFP alone, the fusion protein contains a functional amphipathic helical peptide is able to stimulate a higher level of anti-GFP IgG titer (Figure 1B), whereas the membrane binding defective mutant, AH2M3 (Figure 1C), was unable to activate humoral immunity against GFP. The IgG subtype of the sera collected was then determined using subtype-specific secondary antibodies. The result showed the titer of IgG1 is significantly higher than either IgG2a or IgG2b, an indication that the immune response is tilted toward the Th2 type (Figure 1D).

**Figure 1.**
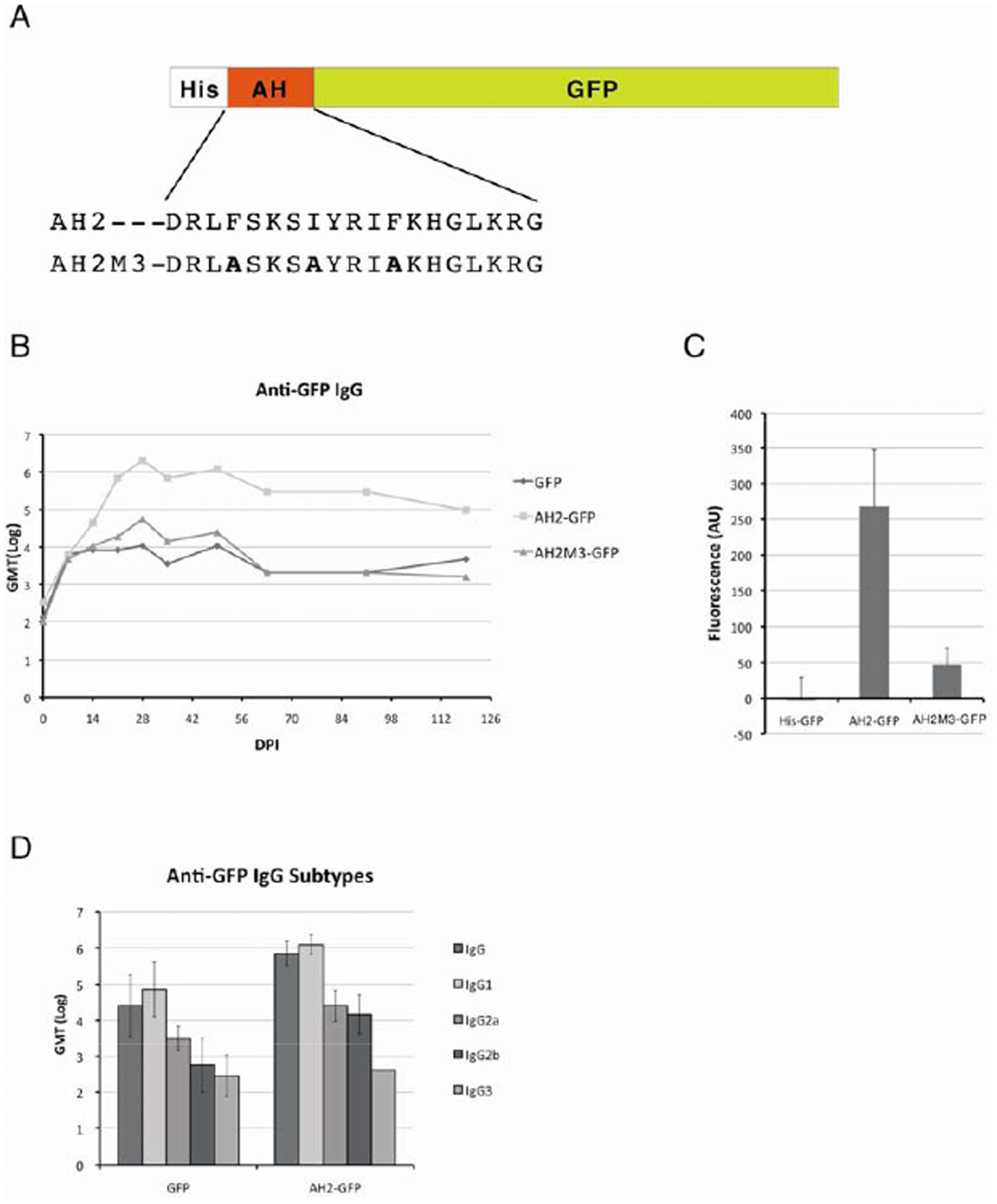
The effect of AH peptide mediated recombinant protein antigenicity enhancement. (A) The design of the AH fusion protein was shown. The AH peptide from the M2 protein of influenza virus strain A/TW/3355/97 H1N1 (AH2) was inserted between His-Tag and GFP. The amino-acid sequence of the AH2 peptide and its membrane binding mutant, AH2M3, were shown. (B) The change of anti-GFP IgG titers was determined by ELISA and followed for 4 months. Mice were immunized twice with 20 μg each of His-GFP, AH2-GFP, or AH2M3-GFP at days 0 and 14 by intramuscular injection. Sera for ELISA assay were collected every 7 days at first month, every 2 weeks up to second month and monthly up to fourth month. The anti-GFP IgG geometric mean titer was shown as GMT (Log) (N=5). (C) The membrane binding activity of fusion proteins was measured by incubating the fusion proteins with cultured MDCK cells for 4 hours and then washing with PBS for quantification under a fluorometer. (D) The anti-GFP IgG subtypes of sera collected at day 28 after the primary dose of His-GFP or AH2-GFP were determined. HRP-conjugated goat anti-mouse IgG1, goat anti-mouse IgG2a, goat anti-mouse IgG2b, and goat anti-mouse IgG3 secondary antibodies were used in the ELISA assay.

One potential explanation of amphipathic helical peptide-mediated immunity boosting is by activating Toll-Like Receptor (TLR) pathways, the major mechanism for boosting recombinant protein antigenicity by adjuvants. To test this hypothesis, we collected the mouse plasma 2 hours after intramuscular injection of either PBS, GFP, AH2-GFP, or TLR5 ligand, flagellin, and examined the expression of inflammatory cytokines, IL-6 and MCP-1. IL-6 and MCP-1 are cytokines that are secreted by macrophages after activation of TLR pathways by invading pathogens or adjuvants. The flagellin is the structural component of bacterial flagellum that controls microbe movement and is the ligand for TLR5. The results showed a dose-dependent induction of IL-6 and MCP-1 expression by injected recombinant flagellin (Figure 2A, 2B). But not when the mice were injected with His-GFP or AH2-GFP recombinant proteins. These results suggest that the activation of humoral immunity by AH2-GFP is independent from the TLR signaling pathways; instead, the activity of AH2-GFP is correlated with its ability to bind plasma membrane.

**Figure 2.**
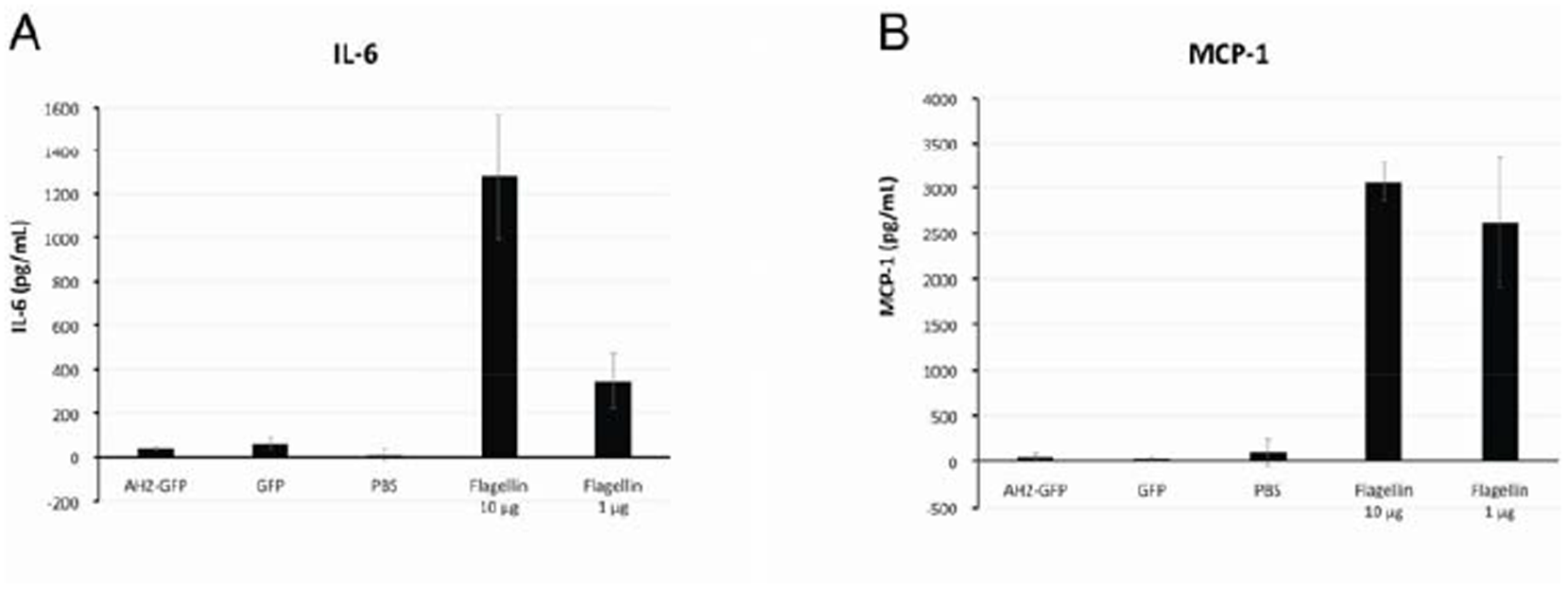
Detection of inflammatory cytokine levels after antigen immunization. Mice were injected with either PBS, His-GFP, AH2-GFP, 10 μg flagellin or 1 μg flagellin and plasma was collected 2 hours post immunization for detection of A) IL-6 and B) MCP-1 by sandwich ELISA. (N=5).

To trace the transportation of AH2-GFP fusion protein after binding to the plasma membrane, we incubated purified AH2-GFP fusion protein with cultured MDCK cells for four hours and evaluated whether AH2-GFP will be endocytosed into the cell. The confocal images showed the endocytosed AH2-GFP in a vesicle-like structure (Figure 3A). To trace the transportation of endocytosed AH2-GFP, we conjugated the AH2 peptide onto the surface of a gold nanoparticle (G-NP) and used a transmission electronic microscope to trace the tansportation of peptide conjugated AH2-G-NP. The cultured cell was either incubated with AH2-G-NP for 4 hours (Figure 3B-C) or 4+20 hours (Figure 3D-E) before being washed and processed for TEM imaging. The results showed that four hours after AH2-G-NP binding to the plasma membrane, AH2-G-NP containing membrane (marked by arrows) was transported to the multivesicular body (MVB), an organelle for membrane protein sorting, or the lysosome (membrane rich and electron dense vesicle, labled L). In 4+20 hours post incubation, many AH2-G-NPs were observed in enlarged lysosomes (L) and many within permeabilized membranes (PL). Some of the enlarged lysosomes were shown with broken membranes, possibly due to the AH2-G-NP and some aggregated AH2-G-NP can be observed being released into the cytoplasm (marked by arrow heads). These results suggest the fusion of GFP with an amphipathic helical peptide not only mediates the binding of fusion protein onto the plasma membrane but also the transport and delivery of fusion protein into the cytoplasm through a permeabilized lysosome.

**Figure 3.**
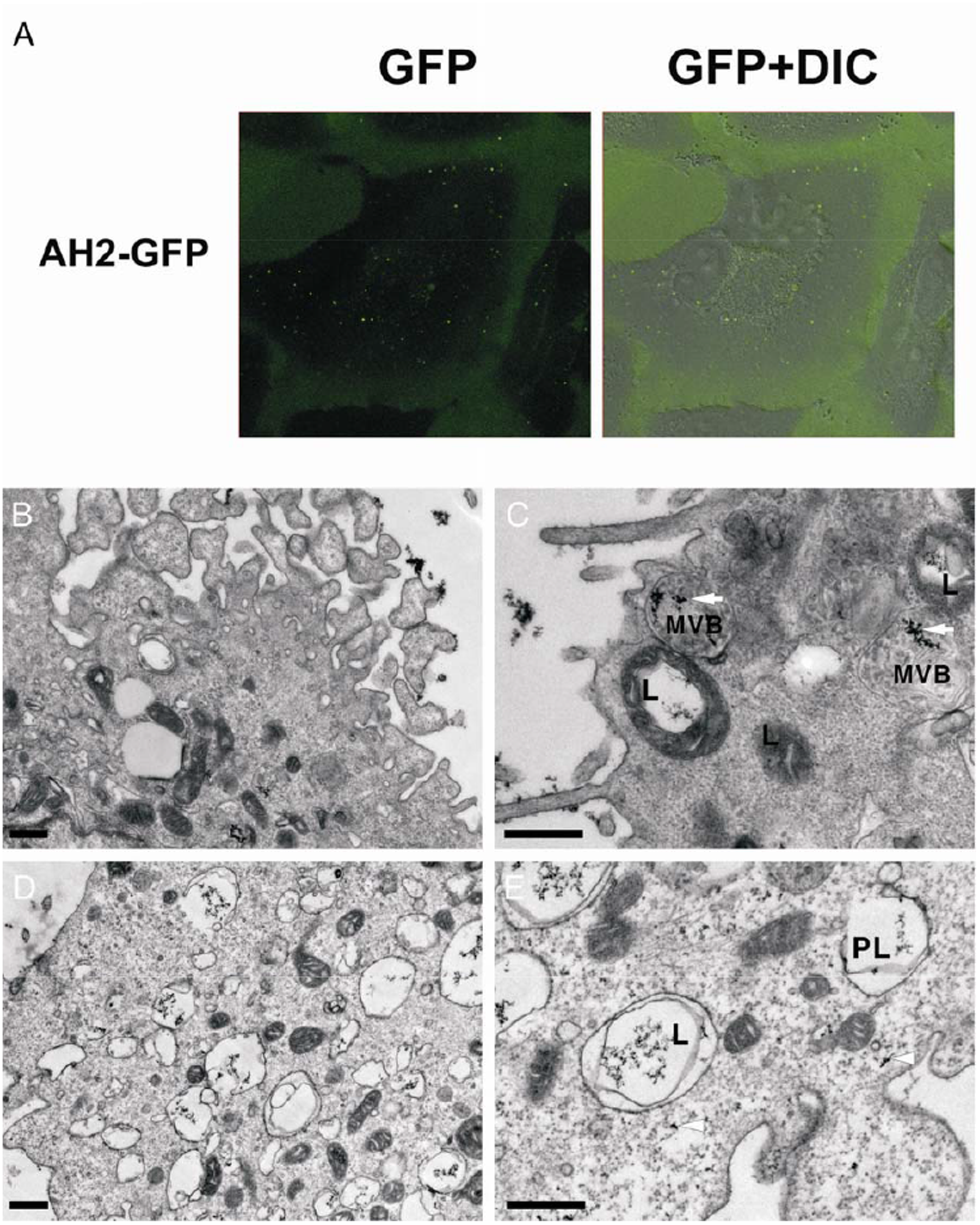
The binding and transportation of AH2-GFP and AH2-G-NP into cytoplasm were examined under a confocal fluorescence microscope or transmission electronic microscope (TEM). (A) Purified AH2-GFP fusion protein was incubated with cultured MDCK cells and non-binding protein was washed away after 4 hours of incubation. The location of internalized AH2-GFP was traced by confocal fluorescence microscopy. The transportation of AH2 peptide conjugated gold nanoparticles (AH2-G-NP) was traced by TEM. The MDCK cell was incubated with AH2-G-NP for either (B-C) 4 hours or (D-E) 4+20 hours before it was fixed and embedded for TEM imaging. The arrow points to the internalized AH2-G-NP in the lysosome (L) or permeabilized lysosome (PL) or the multivesicular body (MVB). The arrowheads pointed to the positions of released AH2-G-NP in cytoplasm. The length of the scale bar is 500 nm.

Amphipathic helical peptides play multiple roles in the protein-membrane interactions, including membrane anchorage, inducing membrane curvature, and membrane lysis [2,8]. To evaluate whether the AH2 peptide possessed membrane lysis activity as implicated in Figure 3C, the synthetic AH2, AH2M3, and Udorn peptides in increasing concentrations were applied to cultured Madin-Darby canine kidney (MDCK) cells. The positive control Udorn peptide, a M2 AH peptide that was derived from the influenza strain A/Udorn/72, has been shown to induce membrane leakage in an artificial liposome that contains low percentage of cholesterol in an in vitro assay [4]. The effect of synthetic peptides on the viability of cultured MDCK cells was determined by flow cytometry for cells undergoing apoptosis and the MTT assay for cells with metabolic activity. The results suggested 20% of the MDCK cells incubated with Udorn peptide became apoptotic at a peptide concentration of 10 μM and the percentage of apoptotic cells increased with elevated peptide concentration (figure 4A). This result is in agreement with a previous study from Lamb’s group [4]. In contrast, the percentage of MDCK cells undergoing apoptosis when treated with AH2 peptide concentrations up to 100 μM was not significantly different from those treated with PBS. In the MTT assay, the mitochondrial dehydrogenase enzyme activity of cells treated with Udorn peptide dropped significantly compared to cells treated with AH2 peptide in all three peptide concentrations. Whereas the viabililty of MDCK cells treated with AH2 peptide only became significantly different from the PBS control when the peptide concentration reached 100 μM. These results suggest the AH2-fusion protein has to reach 100 μM (3.5 mg/ml) to induce a minor reduction in cell viability.

**Figure 4.**
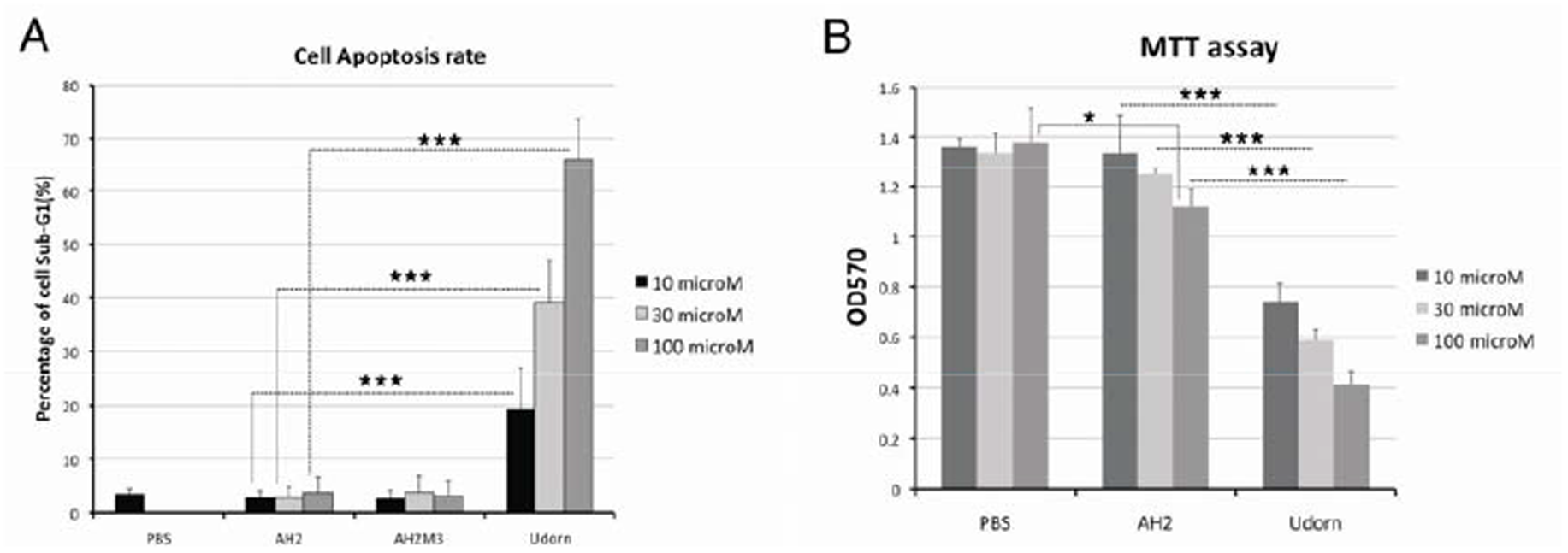
Evaluating the cytotoxic effects of amphipathic helical peptides by flow cytometry and MTT assays. (A) The percentage of cells in apoptosis after being exposed to the AH peptide was measured by flow cytometry. MDCK cells were incubated with 10 microM, 30 microM or 100 microM of synthetic peptides for 24 hrs, and then the cells were harvested for detecting apoptotic cells using nuclei staining. (B) The viability of MDCK cells after being incubated with synthetic peptides for 24 hours was measured by the MTT assay. The AH2 peptide or Udorn peptide was added into the culture media in increasing concentrations of 10 microM, 30 microM or 100 microM. (*** P≤0.001, * P≤0.05)

To test whether AH2-mediated activation of humoral immunity is a particular phenomenum or a common feature of amphipathic helical peptides, we replaced the AH2 peptide with published AH peptides from either NSP1 of the Semliki Forest virus (NSP1), G-protein coupled protein kinase 5 (GRK5), or the fusion peptide of hemagglutinin (HAfp23). These fusion proteins were expressed and purified under the same conditions as AH2-GFP and were used for mouse immunization using a two-dose protocol with a 14-day gap. The results showed all the fusion peptides tested were able to boost the anti-GFP IgG titer by a magnitude about 1000 folds after fusing to GFP. This increase in antibody titer against a recombinant protein is significant when compared to the adjuvants that have been used to boost the antigenicity of subunit vaccines. The use of alum salt increased the anti-GFP IgG titer by 10 fold and the TLR4 ligand MPLA also had similar activities but not as significant as the GRK5 peptide fusion (Figure 5B).

**Figure 5.**
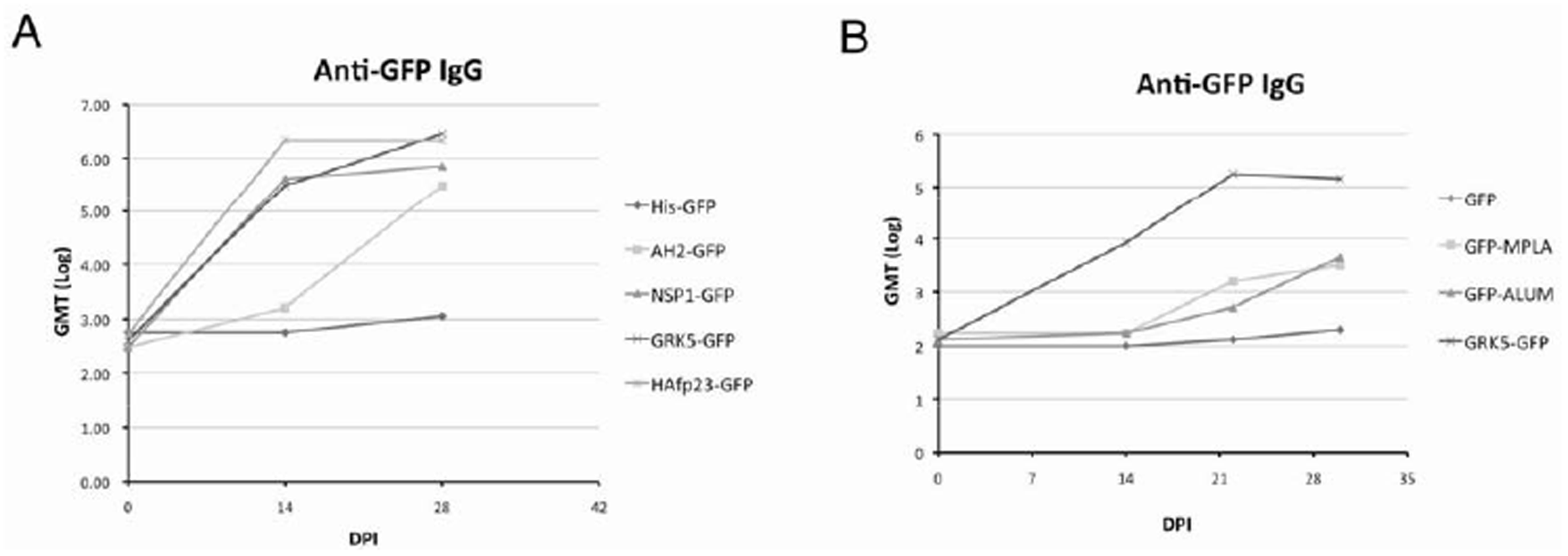
The evaluation of different amphipathic helical peptides in enhancing GFP antigenicity and the comparison with other adjuvants. (A) The AH peptides from M2 protein of H1N1 influenza virus (AH2-GFP), Semliki Forest virus replicase nsP1 (NSP1-GFP), G-protein coupled kinase 5 (GRK5-GFP), and the fusion hairpin peptide of hemaglutinin (HAfp23-GFP) were fused with GFP, and the fusion proteins were purified and used for mouse immunization in a two doses protocol. The anti-GFP IgG titer was determined from sera collected 14 days after immunization. (B) The comparison of GFP antigenicity boosting by either GRK5 peptide (GRK5-GFP), alum salt (GFP-ALUM), or MPLA (GFP-MPLA) in a two-dose immunization protocol. The anti-GFP IgG titer was determined using ELISA. (N=5)

## Discussion

The study results indicate that the AH peptide fusion enhanced the antibody production against the model antigen, GFP. This increase is significant since the comparison with known adjuvants suggests AH peptide fusion is more effective in stimulating antibody production. Adjuvant that activates Th1 type immune responses activates CD4+ T cells and IgG2a expression, whereas adjuvant that activates the Th2 pathway induces IgG1 expression through CD8+ T cells. We have not tested whether the AH peptide fusion also activates cellular immunity against GFP-expressing cells, but the IgG subclass indicates a balanced activation of both T helper type 1 (Th1) and Th2 pathways.

In our results, the binding of AH2-G-NP caused permeabilization of the lysosome membrane but not the plasma membrane. In an in vitro membrane binding study, the M2AH peptide (the same Udorn peptide as in this study) induced membrane leakage in a synthetic liposome when the liposome membrane contained a low percentage cholesterol but not when the membrane contained a high percentage cholesterol [4]. Since the distribution of cholesterol in mammalian cells is organelle dependent, it is rich in the multivesicular body (MVB) and the plasma membrane but low in the lysosome membrane [9], so it is likely that the AH2 peptide only induced membrane leakage when sorted into the lysosome. Another study suggested M2AH binds cholesterol through the aromatic side chains of F47 [5] and the binding of M2AH weakens the lipid bilayer by introducing membrane pits [7]. So the binding of the AH2 peptide likely introduces cholesterol depleted membrane pits and causes lysosomal membrane leakage. This mechanism is similar to that of QS-21, the main component of several clinically approved adjuvants, which induces endocytosis and lysosome pore formation through binding to cholesterol [10]. Other adjuvants that activate immunity through lysosomal leakage include alum salt [11,12].

The AH2-GFP fusion protein immunization did not induce acute expression of IL-6 or MCP-1, two inflammatory cytokines. There are two potential explanations for this result. First, as the QS-21 study suggested, the QS-21 effect on the NF-kB pathway and its downstream effects, including IL-6 expression, is depend on lysosomal maturation, which takes longer than TLR signaling. So the time points (2 hrs post injection) used to determine cytokine level in this study may be too early to catch the IL-6 expression induced by the lysosomal leakage and its downstream NF-κB activation. One study worth mentioning was the delayed induction of IL-6 and other inflammatory cytokines after immunization with alum-salt adjuvanted antigen [13]. Another potential explanation is the distinct pathways that leading to NF-kB signaling and IgG antibody titer induction activated by QS-21 [10]. In this aspect, the AH2 peptide plays a similar role as QS-21, or they share the same ability to destabilize the lysosomal membrane. Applying QS-21 alone causes cell apoptosis, so it is often formulated with cholesterol and phospholipids to reduce cytotoxicity [14-16]. But the formulation of a protein-based vaccine in an oil-in-water nanoemulsion adjuvant may cause instability of the antigen when it is stored at ambient temperature, and it required either lyophilization [17] or spray drying [18] after adding excipients to achieve long-term thermostability. In comparison, protein nanoparticles assembled by an amphipathic helical peptide are self-adjuvanted and can be stored as ready-to-use doses at ambient temperature [19,20].

In this study, several known amphipathic helical peptides were used to evaluate their effects on GFP antigenicity. The GRK5-AH is derived from the C-terminal AH peptide of G protein-coupled protein kinase 5 (GRK5) [21]. Together with the N-terminal helical basic peptide it anchors the GRK5 protein onto the plasma membrane [22]. The NSP1-AH is adopted from the replicase of the Semliki forest virus, nsP1 [23,24]. This peptide is conserved in replicase from all alphaviruses, including Chikungunya virus [25]. The HAfp23-AH is the fusion peptide of hemagglutinin from influenza virus that induces membrane curvature and fusion through two tightly packed hairpin helices buried in the lipid bilayer [26]. There is no similarity in peptide sequences among the four peptides presented in this study or shared mechanism that can fully explain their activity in boosting antigenicity. The common feature is the tendency to interact with phospholipid membranes through their hydrophobic faces. One potential explanation comes from our recent work [19]. When the Cys8 from a modified M2AH peptide, LYRRLE, was replaced by serine, the substitution caused the destabilization and hydrolysis of protein nanoparticles, and the particle size increased from 23 nm to 600 nm, suggesting the particle was a loosely packed aggregation [19]. So the activity of AH peptides in enhancing anti-GFP IgG may reside in their ability to form a random aggregation after fusing with GFP. Other example of GFP fusion proteins forming aggregated megastructures and gaining high antigenicity have been reported when GFP was fused with the N-terminal domain of polyhedrin protein [27]. But the replacement of GFP with known vaccine antigen in this megastructure is difficult because of the interference with protein folding when proteins are packed closely.

Two synthetic peptides were reported to serve the role of an adjuvant. The Q-11 peptide self-assembled into an extended nanofibril structure that stimulated long-lasting humoral and cellular immunity to the covalently fused peptide if proper epitopes were provided [28-30]. The Q-11 peptide’s activities depend on T cells and MyD88, a typical signaling molecule that mediates innate immunity, but no inflammatory responses were observed when immunizing with a Q-11-based vaccine [29]. The other self-assembling peptide is coil-29, which forms an extended spiral tube structure [31]. The coil-29 also shares similar activity with the Q-11, but it has higher potency in inducing both types of immunity [32]. Interestingly, the coil-29 peptide is predicted to be an amphipathic helical peptide according to prediction by HeliQuest [33]. There are several similarities between these two self-assembling peptides and the peptides identified in this study: they are both self-adjuvanting, both show low reactogenicity, and they are all amphipathic peptides, suggesting they might also share similar mechanisms that demand further studies.

In conclusion, the fusion with an amphipathic helical peptide provides a novel mechanism for stimulating immunity and may be applied to other antigens other than GFP if the following goals can be achieved: extended protein stability, efficient incorporation of heterologous proteins, and high specificity to the target cell.

## Materials and methods

### Expression and purification of endotoxin free protein

The pET28a vector encoding the target protein was transformed into BL21 (DE3) using heat shock. The procedure for heat shock transformation is as follows: Day 1 at 3 p.m., plasmid of 1 ng was mixed with 50 μl of thawed competent cell, BL21(DE3), and incubated on ice for 10 minutes before 42 °C heat shock for 40 seconds. After heat shock, the mixture was kept on ice for 1 minute before adding 1 ml LB and then shaken at 220 rpm at 37 °C for 1 hour. Then the bacteria were spun down at 13000 rpm for 1 minute. The bacteria pellet was re-suspended in 100 μl LB and plated on LB plates with 100 μg/ml ampicillin and incubated at 37 °C. Day 2, at 10 a.m., there should be fewer than 100 colonies on the plate. Pick one colony and inoculate the bacteris in 3 ml LB/ampicillin culture and incubated by shaking at 37(/225 rpm. By 12 a.m., the 3 ml culture was transferred to 500 ml LB/ampicillin media, and the growth of culture was monitored by spectrophotometer. When OD600 reached 0.5-0.7, 1 mM IPTG was added into the culture to induce protein expression. The culture was harvested 3 hours after protein induction. Bacteria were spun down by centrifuge at 5000 rpm for 10 minutes. Supernatant were removed and the pellet was kept for further processing.

### Protein purification

The buffer for protein purification and gel filtration: 20 mM NaPO_4_, pH 7.4, and 300 mM NaCl (GF buffer). For Ni-NTA resin purification: 5 mM Imidazole in GF for bacteria lysis (Lysis buffer), 10 mM Imidazole in GF for wash (Wash buffer), 200 mM Imidazole in GF for protein elution (Elution buffer). Bacteria pellets from 500 ml of LB culture was re-suspended in 40 ml of Lysis buffer by vortex and then was lysed using an ultrasonic sonicator (Misonix 3000) at 10 S on/5 S off cycles for 5 minutes. The re-suspended bacteria were kept in icy water during sonication. Insoluble debris was removed by centrifugation at 10000 rpm for 10 minutes using a Sorval SS34 rotor at 4 °C. The supernatant containing target proteins was passed through a polypropylene column containing 2 ml Ni-NTA resin from Qiagen. After the target protein bound to the Ni-NTA resin, the column was washed with 10 bed volume of wash buffer (20 ml). After washing, the target protein was then eluted using 5 ml elution buffer. Target protein purified using Ni-NTA resin was further purified using a gel filtration column from GE (HiLoad 16/600 Superdex 200 pg) on AKTA prime.

### Size exclusion chromatography

Three programs were setup for the purification of fusion proteins in study. Program 30: For gel filtration purification of the fusion protein. Program 33: for the onsite cleaning of HiLoad 16/600 Superdex 200 pg using 120 ml of 0.5 M NaOH. The flow rate is 0.8 ml/minute. Program 34: for equilibrate the column after onsite cleaning using 150 ml GF buffer. The column was washed after two protein purifications. A 5 ml sample loop was connected to the machine, the AKTA Prime, for holding the protein sample. The sample loop was first flushed with 25 ml of GF buffer to remove the residual protein from previous purification. Before loading the fusion protein into the sample loop, additional 0.5 ml of GF buffer was added into the Ni-NTA eluted fusion protein. The protein solution was loaded into a 5-ml syringe and injected into the sample loop. The glass tubes in the corresponding fractions of fusion protein eluted from the column were replaced with new tubes every time before the purification. All the buffers to be used in the gel filtration experiment were filtered through a 0.2-μm filter unit for sterilization and impurity. These buffers were further degassed by vacuum for 10 minutes. The FPLC fractions contained fusion proteins as monitored by UV280 and were collected and pooled. The pooled protein solution was concentrated by a second Ni-NTA column with the same bed volume as the first one. After the protein was eluted from 2^nd^ Ni-NTA column, the protein concentration was determined by BCA analysis. Two fractions with highest protein concentration were pooled and dialyzed against 1 L PBS overnight. After dialysis, the endotoxin contaminant was removed using EndotoxinOUT™ resin from G-Bioscience. The polypropylene column was filled with 4 ml EndotoxinOUT™ resin and washed with 5 bed volumes of 1 % sodium deoxycholate in LAL Reagent Water (LRW), 5 bed volumes of LRW and 5 bed volumes of PBS/LRW before loading 1.5 ml fusion protein. After complete loading of the protein, the column was placed in cold room for 1 hour for complete absorption of endotoxin by resin. Protein was eluted using stepwise additions of first fraction of 0.5 ml and later fractions of 1 ml of PBS.

### LAL assay

The endotoxin level was determined using PYROTELL-T LAL provided by Associate of Cape Cod. Each LAL reagent supplied in lyophilized powder was first dissolved in 5 ml LRW and then aliquoted into small volumes and kept at -80 °C. In a typical turbidity assay using PYROTELL-T, 100 μl of sample, including endotoxin standard, were applied to a microplate. The reaction kinetics were measured using a victor3 fluorescence reader by OD405 at 37 °C. The time-lapse between each reading was 70 seconds and the reading was repeated 60 times. The time needed for a reaction to reach a difference of 0.1 in OD405 between sample and blank was recorded and compared to the endotoxin standard to determine the endotoxin unit in a sample. The log of endotoxin activity was plotted against time (seconds) to reach a difference of 0.1 in OD405; a logarithmic trend line is used as the standard curve. The identified endotoxin activity in a protein sample was further combined with the protein concentration, as determined by BCA assay, to determine specific endotoxin activity, EU/mg. The upper limit of specific endotoxin activity in all the protein samples was 5 EU/mg.

### Animal Immunization

The C57BL/6 SPF mice used in immunization procedures were between aged 6 and 8 weeks and provided by BioLasco Taiwan. Without anesthetization, mouse was bled from tail by clipping away 2-3 mm tail. Blood was collected using a pipette with occasional massaging on the tail. About 100 μl blood was collected from each bleed. Blood was left at RT for 2 hours before centrifugation at 2000 rpm for 10 minutes in a microcentrifuge, and serum was collected in a microtube. Remaining lymphocytes were removed by centrifugation at 2000 rpm for 10 minutes in a microcentrifuge. Serum was stored at -80 °C before use in an ELISA assay. Mouse tail bleeding is stopped by heated soldering iron while pressing the tail tip between fingers. For primary immunization, 20 μg purified protein was injected intramuscularly in the thigh of the hind limb. Blood was bled 14 days post immunization from tail and a booster dose with the same amount of purified protein was injected by the same procedure.

### ELISA for determining antibody titer

The anti-GFP IgG titer was determined by coating GFP protein at a concentration of 10 μg/ml. The procedures for the ELISA assay followed the standard protocol. Secondary Ab (HRP conjugated goat anti-mouse Ab) was diluted by 5000 folds in blocking buffer. The coating buffer contains 0.2 M Carbonate/Bicarbonate buffer at pH 9.4. The washing buffer contains 0.05% Tween 20 in 1X PBS. The blocking buffer contains 1% BSA in the washing buffer, and the stop buffer contains 2M sulfuric acid.

### GFP fusion protein binding assay

The GFP fusion protein binding activity was tested by first plating 4×10^4^ MDCK cells on one well of a 96 well plate, each sample was tested in quadruplicate. On second day, 200 μg/ml of GFP fusion protein was added into the culture media and incubated for 4 hours. After incubation, media was removed and washed with 200 μl of PBS for three times. The amount of GFP fusion protein bound/internalized was determined by a fluorometer.

### Plasma cytokine determination

To detect the blood cytokine level, mice were first intramuscularly injected with experimental proteins. Two hours after injection, the mice were anesthetized before collecting blood through cardiac puncture with a syringe preloaded with 10 μl of 0.5M Na_2_EDTA. The blood samples were centrifuged at 2000 rpm for 20 minutes in a tabletop microcentrifuge. The plasma collected was aliquoted into 20 μl fractions and snap-frozen on dry ice. The plasma cytokine level was determined by sandwich ELISA with a standard from eBioscience, USA. The IL-6 level was detected using the Mouse IL-6 ELISA Ready-SET-Go! (Cat#88-7064, eBioscience, USA). The MCP-1 level was detected using the Mouse CCL2 (MCP-1) ELISA Ready-SET-Go! Kit [34-7391, eBioscience, USA]. The detection procedures followed the manufacturer’s standard protocol.

### Cell apoptosis assay by flow cytometer and MTT assay

MDCK cells were first plated on 6 well plates at 3×10E5 cells/well. Twenty-four hours after plating, peptides were applied to 10 μM, 30 μM, or 100 μM concentrations and cells were incubated for 24 hours before harvested by trypsin digestion. The suspended cells due to apoptosis were collected by centrifuging the culture medium at 1000 rpm for 5 minutes. The cells collected from trypsin digestion and cultured media were combined, and cells were fixed in 3 ml 70 % ethanol overnight in a -20 °C freezer. After fixation, cells were washed and resuspended in 1 ml PBS and stained by the SYTOX AADvanced Dead Cell Stain Kit (Thermo Fisher Scientific Inc.) in the presence of 100 μg RNaseA for 30 minutes at 4 degree cold room before analysis by flow cytometry (Calibur BD). The sub-G1 fraction was calculated and designated as apoptotic cells. For MTT assay, MDCK cells were plated at 1×10E4 cell/well. Twenty-four hours after plating, peptide was added to reach a concentration of 10 μM, 30 μM, or 100 μM and cells were incubated for 24 hours and media removed and replaced with 50 μl serum free media and 50 μl MTT solution. After incubation at 37 °C for 3 hours, 150 MTT solvent was added and the plate was shaked for 15 minutes and then read the absorbance at OD570.

### Transmission electronic microscope

Madin-Darby canine kidney (MDCK) cells were seeded on ACLAR EMBEDDING film in a 6-well plate at a density of 3×10^5^ cells per well. After overnight culturing, part of the media were removed to leave 0.5 ml of culture media before adding 50 μl of 10 nm gold nanoparticles conjugated with AH2 peptide. The peptide density of AH2 conjugated 10 nm gold nanoparticle used in this assay was 4.78×10^12^/ml. After 4 hours of incubation, cells were washed in serum-free culture media at room temperature and fixed directly or cultured for an additional 20 hrs before fixation. The fixation procedure involved covering the film in 2.5% glutaraldehyde, 0.1% tannic acid, 0.1M cacodylate buffer, pH 7.2 at room temperature for 30 minutes. Then cells were washed in 0.1M cacodylate buffer pH 7.2, 0.2M sucrose, 0.1% CaCl_2_ for 5 minutes at room temperature. Afterward, cells were post-fixed in 1% OsO_4_/0.1M cacodylate buffer, pH 7.2 at room temperature for 30 minutes. Then cells were washed with ddH_2_O three times for 5 minutes each at room temperature. Cells were further stained with 1% uranyl acetate at room temperature for 30 minutes before being washed in ddH_2_O for 3 times. Cell were then dehydrated and embedded in EPON by polymerizing at 60 °C for 24 hours. Embedded cells were then sectioned for observation under a transmission electronic microscope.

